# Harmonization of Structural Brain Connectivity through Distribution Matching

**DOI:** 10.1101/2024.09.05.611489

**Authors:** Zhen Zhou, Bruce Fischl, Iman Aganj, the Alzheimer’s Disease Neuroimaging Initiative

**Author notes:** Corresponding authors {, }. Data used in the preparation of this article were obtained from the Alzheimer’s Disease Neuroimaging Initiative (ADNI) database (adni.loni.usc.edu). As such, the investigators within the ADNI contributed to the design and implementation of ADNI and/or provided data but did not participate in the analysis or writing of this report. A complete listing of ADNI investigators can be found at http://adni.loni.usc.edu/wp-content/uploads/how_to_apply/ADNI_Acknowledgement_List.pdf.

## Abstract

The increasing prevalence of multi-site diffusion-weighted magnetic resonance imaging (dMRI) studies potentially offers enhanced statistical power to investigate brain structure. However, these studies face challenges due to variations in scanner hardware and acquisition protocols. While several methods for dMRI data harmonization exist, few specifically address structural brain connectivity. We introduce a new distribution-matching approach to harmonizing structural brain connectivity across different sites and scanners. We evaluate our method using structural brain connectivity data from three distinct datasets (OASIS-3, ADNI-2, and PREVENT-AD), comparing its performance to the widely used ComBat method and the more recent CovBat approach. We examine the impact of harmonization on the correlation of brain connectivity with the Mini-Mental State Examination score and age. Our results demonstrate that our distribution-matching technique effectively harmonizes structural brain connectivity while maintaining non-negativity of the connectivity values, and produces correlation strengths and significance levels competitive with alternative approaches. Qualitative assessments illustrate the desired distributional alignment across datasets, while quantitative evaluations confirm competitive performance. This work contributes to the growing field of dMRI harmonization, potentially improving the reliability and comparability of structural connectivity studies that combine data from different sources in neuroscientific and clinical research.

## 1 Introduction

Diffusion-weighted magnetic resonance imaging (dMRI) is a powerful noninvasive technique for probing the microstructure of biological tissue, particularly the brain white matter (Tournier et al., 2011). By measuring the diffusion of water molecules, dMRI allows us to infer the organization and integrity of neural tissue, making it instrumental in both neuroscientific research and clinical applications. Structural brain connectivity (Bazinet et al., 2023), typically represented as networks derived from dMRI fiber tracking (tractography) (Behrens et al., 2007; Mori et al., 1999), provides crucial insights into the architecture of white-matter pathways and the organization of communication pathways in the brain (Bazinet et al., 2023; Yeh et al., 2021).

dMRI has been widely used to study brain development, aging, as well as various neurological and psychiatric disorders (Beck et al., 2021; Frau-Pascual et al., 2021; Pines et al., 2020; Wheeler & Voineskos, 2014), with recent years seeing an increase in multi-site dMRI studies to investigate brain disorders on a larger scale, such as the Alzheimer’s Disease Neuroimaging Initiative (ADNI) (Jack Jr et al., 2015; Weiner et al., 2015), the Parkinson’s Progression Markers Initiative (Marek et al., 2011), and in Huntington’s disease research (Magnotta et al., 2012). These studies offer the potential for increased statistical power and the ability to detect subtle effects that may not be apparent in smaller, single-site studies. However, the variability in scanner hardware, acquisition protocols, and processing methods across different sites can introduce unwanted variability in the data, potentially confounding biological effects of interest. This challenge has highlighted the critical need for robust methods to harmonize dMRI data between different scanners, protocols, and populations (Zhu et al., 2019).

Several approaches have been proposed to address the harmonization problem. One widely used method is ComBat, originally designed for genomic data and later adapted for neuroimaging (Johnson et al., 2007). ComBat and its variants have been successfully applied to a variety of neuroimaging studies to harmonize diffusion tensor imaging (DTI) measures, cortical thickness, regional volumetric measures, and functional connectivity properties (Fortin et al., 2017, 2018; Pomponio et al., 2020; Yu et al., 2018; Zhou et al., 2022, 2023). More recently, an extension of ComBat, called CovBat, was developed to mitigate site effects in covariance for machine learning in neuroimaging data (Chen et al., 2022). However, a limitation of ComBat and related methods is that its optimization procedure assumes a normal distribution for the data, which may not necessarily be appropriate for all types of neuroimaging measures or data (Johnson et al., 2007; Pinto et al., 2020; Pomponio et al., 2020).

Another notable method designed explicitly for dMRI data harmonization is the Rotation Invariant Spherical Harmonics (RISH) approach, introduced by Mirzaalian et al. (2016). RISH features are computed from spherical harmonic decompositions of the diffusion signal, with linear mappings used to harmonize these features across sites. Karayumak et al. (2019) extended this method for varying acquisition parameters, whereas De Luca et al. (2022) validated it on multi-shell dMRI data from three sites. Building on RISH, several deep-learning methods have been developed to leverage Convolutional Neural Networks to harmonize dMRI data (Koppers et al., 2017, 2019; Tax et al., 2019), preserving rotational invariance while leveraging the power of deep-learning architectures to learn complex mappings between scanner-specific features. More recently, an adaptive template estimation approach combined with RISH features was proposed to better account for inter-subject anatomical variability when harmonizing voxel-wise dMRI signals or surface-based features across different scanners (Xia & Shi, 2022, 2024). The RISH method requires well-matched control subjects (16-20 per site) to learn harmonization parameters, which can be challenging for retrospective or rare-condition studies. Furthermore, harmonizing diffusion images early in the pipeline necessitates rerunning subsequent processing steps, which increases the computational time.

A few other notable approaches to dMRI harmonization are as follows. The Method of Moments (MoM), proposed by Huynh et al. (2019), matches the first and second moments (spherical mean and variance) of the diffusion signal across sites using a linear mapping function. To capture all sources of scanner-specific variations, Zhang et al. (2023) proposed RELIEF (REmoval of Latent Inter-scanner Effects through Factorization), which uses a structured multivariate approach based on linked matrix factorization to model and remove sources of scanner effects such as scanner-specific means, variances, and latent patterns. Moyer et al. (2020) proposed an unsupervised approach using variational autoencoders to learn scanner-invariant representations, which can map dMRI data between different scanners and protocols without requiring paired training data, while preserving biologically meaningful information.

While these methods have shown success in harmonizing the diffusion signal and derived measures such as fractional anisotropy and mean diffusivity, fewer approaches have been developed explicitly for harmonizing structural brain connectivity (a.k.a. the connectome). Importantly, structural connectivity may be particularly sensitive to scanner and protocol differences (Panman et al., 2019; Zhu et al., 2019), as well as the parameters specified for parcellation and tractography (Sotiropoulos & Zalesky, 2019; Yeh et al., 2021), highlighting the need for robust harmonization methods specifically tailored to the quantified structural connectivity (Kurokawa et al., 2021; Onicas et al., 2022; Patel et al., 2024). To address the gap in harmonization methods for structural connectivity, we propose a new approach, based on distribution matching (DM), to harmonize structural brain connectivity across different sites and scanners. The DM method aligns the statistical properties of different datasets to minimize biases and improve comparability (Bishop & Nasrabadi, 2006; Evans & Rosenthal, 2004). This process aligns the data distributions using methods ranging from simple linear transformations to complex techniques such as quantile normalization or histogram matching. By reducing dataset-specific variabilities, DM enhances the reliability of subsequent analyses when integrating data from various sources. In a previous study, structural MR images were harmonized across different scanners and sites by aligning voxel intensity distributions (Wrobel et al., 2020), demonstrating improved reduction of scanner-related variability while preserving biological differences in multi-site neuroimaging studies compared to existing techniques. In this study, we validated our method by assessing its performance in harmonizing multi-site structural brain connectivity data while comparing it with the ComBat and CovBat approaches.

The rest of the paper is organized as follows. Section 2 introduces the three datasets, the DM approach, and the corresponding implementation. Experimental results are presented in Section 3 to qualitatively and quantitatively demonstrate the ability of the proposed method to harmonize structural brain connectivity in comparison with other methods. Finally, discussions and conclusions are provided in Section 4.

## 2 Materials and methods

### 2.1 Datasets

We used the following three public dMRI datasets in our analysis, with the number of subjects indicating those processed and included in our study (see Table 1): the third phase of the Open Access Series of Imaging Studies (OASIS-3) (LaMontagne et al., 2019) comprising 761 cognitively normal and AD subjects, the second phase of the Alzheimer’s Disease Neuroimaging Initiative (ADNI-2) (Beckett et al., 2015) including 209 participants ranging from cognitively normal individuals to those diagnosed with AD, and the Pre-symptomatic Evaluation of Experimental or Novel Treatments for Alzheimer’s Disease (PREVENT-AD) dataset with 340 subjects at risk of AD (Tremblay-Mercier et al., 2021). Compared to a previous study where we used these datasets (Aganj et al., 2023), here we excluded 10 and 8 outlier subjects from OASIS-3 and ADNI-2, respectively, during quality check. We included the Mini-Mental State Examination (MMSE) score (available in OASIS-3 and ADNI-2) and age (available in all three datasets) as non-imaging variables in our correlation analysis.

**Table 1:** Demographic information and scanner types of the OASIS-3, ADNI-2, and PREVENT-AD datasets.

| Dataset | OASIS-3 | ADNI-2 | PREVENT-AD |
| --- | --- | --- | --- |
| Number of subjects | 761 | 209 | 340 |
| Percentage of males | 44.0% | 55.5% | 28.5% |
| Age (mean ± std) | 69.4 ± 9.0 | 73.5 ± 6.9 | 63.2 ± 5.0 |
| MMSE (mean ± std) | 28.2 ± 2.7 | 27.2 ± 2.6 | — |
| Scanner | Siemens<br>(Trio Tim & Biograph mMR) | GE | Siemens<br>Trio Tim |

We found significant correlations between some of the demographic/clinical variables within these databases, namely between age and sex (OASIS-3: *r* = 0.14, *p* = 0.0002, age was higher for male), between age and MMSE (OASIS-3: *r* = −0.28, *p* = 0; ADNI-2: *r* = −0.14, *p* = 0.048), and between sex and MMSE (OASIS-3: *r* = −0.13, *p* = 0.0004, MMSE was higher for female).

### 2.2 Data processing

Anatomical MR images from the three datasets were processed using FreeSurfer (Fischl, 2012). We included each subject only once, specifically the earliest visit containing dMRI (which was often the baseline visit), to maintain the independence of our data points through a cross-sectional study design.

We then ran the FreeSurfer dMRI processing pipeline, which also incorporates commands from the FMRIB Software Library (FSL) (Jenkinson et al., 2012). This process involved propagating 85 automatically segmented cortical and subcortical regions from the structural to the diffusion space using boundary-based image registration (Greve & Fischl, 2009).

Next, we used our publicly available toolbox (www.nitrc.org/projects/csaodf-hough) to reconstruct the diffusion orientation distribution function in constant solid angle (CSA-ODF) (Aganj et al., 2010) and run Hough-transform global probabilistic tractography (Aganj et al., 2011) to generate an optimal (highest-score) streamline passing through each of the 10,000 seed points for each subject. We computed a symmetric structural connectivity matrix with non-negative elements for each subject by summing the tracts passing through each pair of ROIs weighted by the tract score, which is a function of the ODFs and generalized fractional anisotropy. We then augmented the matrices with indirect connections using the mathematical framework for electrical conductance calculation (Aganj et al., 2014), resulting in a new matrix that additionally reflects multi-synaptic pathways. More details are provided in our previous study (Aganj et al., 2023).

We transformed the connectivity values *c* (each element in the connectivity matrix) into the logarithmic space as *c ←* log(1 + *c*). This transformation helps to reduce the impact of outliers, enabling more robust distribution estimation.

### 2.3 Distribution matching (DM)

The goal of DM is to align the statistical properties of different datasets, ensuring that they share similar distributions. The positive values in our structural connectivity matrices are lower-bounded by zero, peak at some positive value, and have a tail with no theoretical upper bound. This led us to choose the gamma distribution as the appropriate model to represent our (nonzero) connectivity data, as it is well suited for modeling positive continuous data with a skewed distribution, which is typical for structural connectivity values. Characterized by its shape and scale parameters, the gamma distribution is formulated as:

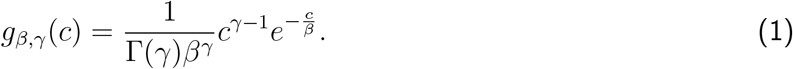

In our case, *c ≥* 0 is the structural connectivity value, *β >* 0 scales *c* hence named the scale parameter, *γ ≥* 1 is the shape parameter, and Γ is the gamma function (Γ(*γ*) = (*γ −* 1)! for integer *γ*). Note that the distribution domain has a lower bound of zero, but is unbounded in the positive direction. Raw (original unaugmented) structural connectivity matrices often contain numerous zero values, indicating the absence of direct connections between certain brain regions. Accordingly, we formulated the probability density function (PDF) of each element in our original (unaugmented) structural connectivity matrix as a combination of an asymmetric variation of the Dirac delta function at the origin and the gamma function, as follows:

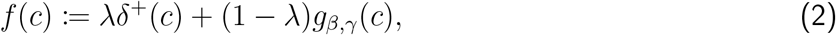

with the following asymmetric variation of the delta function that complies with the nonnegativity of brain connectivity:

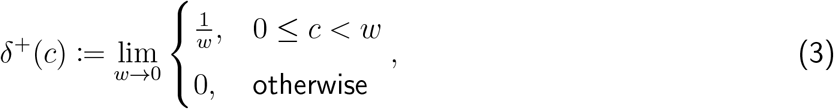

where 0 *≤ λ ≤* 1 represents the portion of zeros in a connectivity value across all subjects. One can verify that *f* (*c*) is normalized.

The above PDF describes the likelihood of the continuous random variable (structural connectivity) taking on the specific value *c*. The cumulative distribution function (CDF), 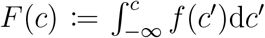, on the other hand, represents the probability of connectivity taking a value no larger than *c*. By applying the *inverse* CDF, also known as the quantile function, we transform the data from one distribution to match another, thereby achieving distributional alignment.

Given two datasets, collected at a reference site R and a new site N, with CDFs *F*_*R*_ and *F*_*N*_, respectively, we want to calculate the transformation *T*_*N*→*R*_ that takes a connectivity value from the new site, *c*_*N*_, and returns a value harmonized in the reference site, *c*_*R*_ = *T*_*N*→*R*_(*c*_*N*_). The harmonization task is to find the value *c*_*R*_ that shares the same percentile in the reference site as that of *c*_*N*_ in the new site, which is accomplished by subsequently applying the CDF of the new site and then the inverse CDF of the reference site to the connectivity value, i.e.,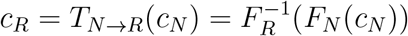, or:

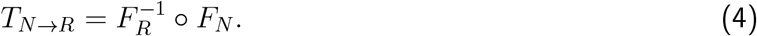

For our gamma+delta distribution model, with the PDF in Eq. (2), the CDF is calculated as:

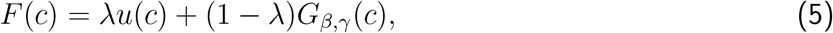

where *u* is a variation of the Heaviside step function (*u*(*c*) is 0 for *c ≤* 0 and 1 otherwise) and *G*_*β*,*γ*_ is the CDF corresponding to *g*_*β*,*γ*_. The inverse of the CDF in Eq. (5), which is used in Eq. (4), has the following closed form:

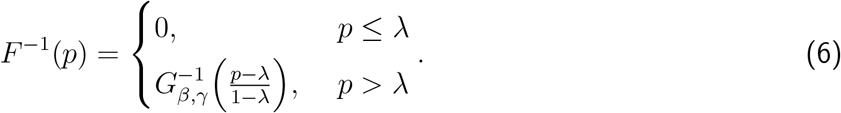

Applying *T*_*N*→*R*_ to a connectivity value from the new site harmonizes it into a value that preserves its percentile in the distribution of the reference site, thereby achieving distributional alignment.

### 2.4 Experimental design

To implement our DM approach, we need to estimate the parameters *λ, β*, and *γ* for each brain connection in both the reference and new sites. For each brain connection across subjects within a site, we first determine *λ* as the empirical portion of zero values in the data. This direct estimation of *λ* represents the probability mass at zero for that particular connection. Note that zero connectivity values in the new site are always mapped to zero after harmonization. Some nonzero values in the new site may also be mapped to zero after harmonization if the new site has a smaller portion of zero values (*λ*) than the reference site.

For the parameters of the gamma distribution (*β* and *γ*), we employ maximum likelihood estimation using MATLAB’s *gamfit* function on the nonzero connectivity values. This parameter estimation procedure is performed independently for each brain connection in each site. For the reference site, these parameters define the target distribution to which the new site’s data will be aligned. When performing harmonization, we estimate these parameters once for each site, and then apply the transformation defined in Eq. (4) to harmonize the connectivity values of the new site to match the distribution of the reference site.

We implemented two variations of our DM approach, so as to account for potential sex differences in structural connectivity: one where we performed harmonization by separating male and female subjects into distinct groups (“DM w/ sex”), and another where we applied harmonization to all subjects together without sex separation (“DM w/o sex”). When separating by sex, we estimated distribution parameters and applied harmonization transformations for male and female groups independently, which allows the method to account for sex-specific connectivity patterns that might otherwise be obscured in a combined analysis.

We used OASIS-3 and PREVENT-AD as reference sites in separate DM experiments. For each referencesite configuration, we tested both sex-separated and non-separated approaches. Analyzing the brain connections between each pair of the 85 cortical and subcortical regions results in 3570 unique connectivity values. We harmonized each original connectivity value (across all subjects) independently and then re-augmented the harmonized values (see Section 2.2). After harmonization, we undid the log transformation as *c ← e*^*c*^ − 1 to ensure the interpretability of the results.

In addition, we performed internal harmonization within the OASIS-3 dataset that includes data from two types of Siemens MRI scanners (655 subjects scanned by Trio Tim and 106 subjects by Biograph mMR). For internal harmonization, we considered subjects scanned by Trio Tim as the reference site while the others were treated as the new site, and applied both sex-separated and non-separated approaches.

In all experiments, for comparison, we also harmonized the structural brain connectivity data using variations of ComBat and CovBat. For each of ComBat and CovBat, we tested two variations: (1) including sex (but not age) as a covariate, and (2) without including either age or sex as a covariate. (See Section 4 for why we did not include age as a covariate in any method.) In contrast to the proposed DM, neither ComBat nor CovBat preserves the non-negativity of the connectivity values, which is required for augmentation. Consequently, to harmonize augmented connectivity with these methods, we applied them directly to augmented connectivity (rather than applying them to original connectivity and re-augmenting the results, as we did with DM). We systematically compared the results of all harmonization approaches with those obtained from our DM method, evaluating the choice of method as well as the impact of reference-site selection (when using our approach).

### 2.5 Correlation analysis

We used MMSE scores for subjects in OASIS-3 and ADNI-2, and age for subjects across all three datasets (OASIS-3, ADNI-2, and PREVENT-AD) to investigate how these variables correlated with structural brain connectivity before and after harmonization and determine if harmonization improved the outcomes of correlation analysis. For robustness, we computed the correlation on a transformed version of the connectivity value (each element in the original or augmented connectivity matrix) as 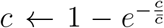, where 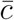 is the cross-subject average of *c*, thereby confining the connectivity values to the range [0, 1]. We started by performing Pearson’s correlation analysis to correlate original (*c*_*O*_) and augmented (*c*_*A*_) connectivity values with MMSE within OASIS-3 and ADNI-2 individually, and with age within each of the three datasets individually.

In our correlation analyses, we examined the proposed DM as well as ComBat and CovBat, all both with and without controlling for sex. For age correlation analyses, we performed the DM experiments twice, each time with a different reference site (OASIS-3 or PREVENT-AD).

We then applied one-sided Wilcoxon signed-rank tests to the absolute values of the correlation coefficients between MMSE/age and structural brain connectivity to measure the effects of the harmonization methods on the correlation strength. These tests were conducted once for each harmonization method against no harmonization. We used a total of four measures to assess the performance: the absolute value of the correlation coefficient (averaged across connections), the *p*-value generated by the abovementioned one-sided Wilcoxon signed-rank test, the number of brain connections surviving the Bonferroni correction (i.e., multiplication of the *p*-value by the number of connections, 3570), and the lowest Bonferroni-corrected *p*-value generated from the correlation analysis across brain connections.

### 2.6 Permutation within ADNI-2

To investigate the potential impact of the reduction in the number of subjects during internal harmonization, we conducted a permutation analysis focusing on the two brain connections most strongly correlated with age and MMSE. We selected a random subset of 106 subjects from the 209 subjects of the ADNI-2 dataset, mirroring the reduction in the number of subjects in our internal harmonization experiment within OASIS-3, and estimated the distribution of the selected connectivity value. We repeated this random selection 1000 times to create a robust permutation set, which helped to elucidate the potential variability introduced by sample size reduction relative to the gold-standard distribution obtained from the full set.

## 3 Results

### 3.1 Qualitative assessment of distribution matching

We measured how much the significance values (*s* := − log *p*), corresponding to the correlation between MMSE and structural brain connectivity, increased after DM harmonization, i.e. Δ*s*, for all brain connections. We then sorted the connections with respect to their Δ*s* and chose the five connections with the smallest, 25^*th*^ percentile, median, 75^*th*^ percentile, and largest Δ*s*. As shown in Fig. 1, the peaks of the fitted gamma curves for ADNI-2 and OASIS-3 better align after (than before) harmonization. The figure also includes results by the ComBat and CovBat harmonization methods (fourth and fifth rows). As illustrated in Fig. 1, ComBat and CovBat, as opposed to our DM approach, may transform zero values into non-zero values after harmonization, creating irregular, multimodal distributions with distinct peaks near zero in several connectivity features. ComBat and CovBat also transform some connectivity values to the implausible negative region, which we projected to zero in the figure to improve visualization.

**Figure 1:**
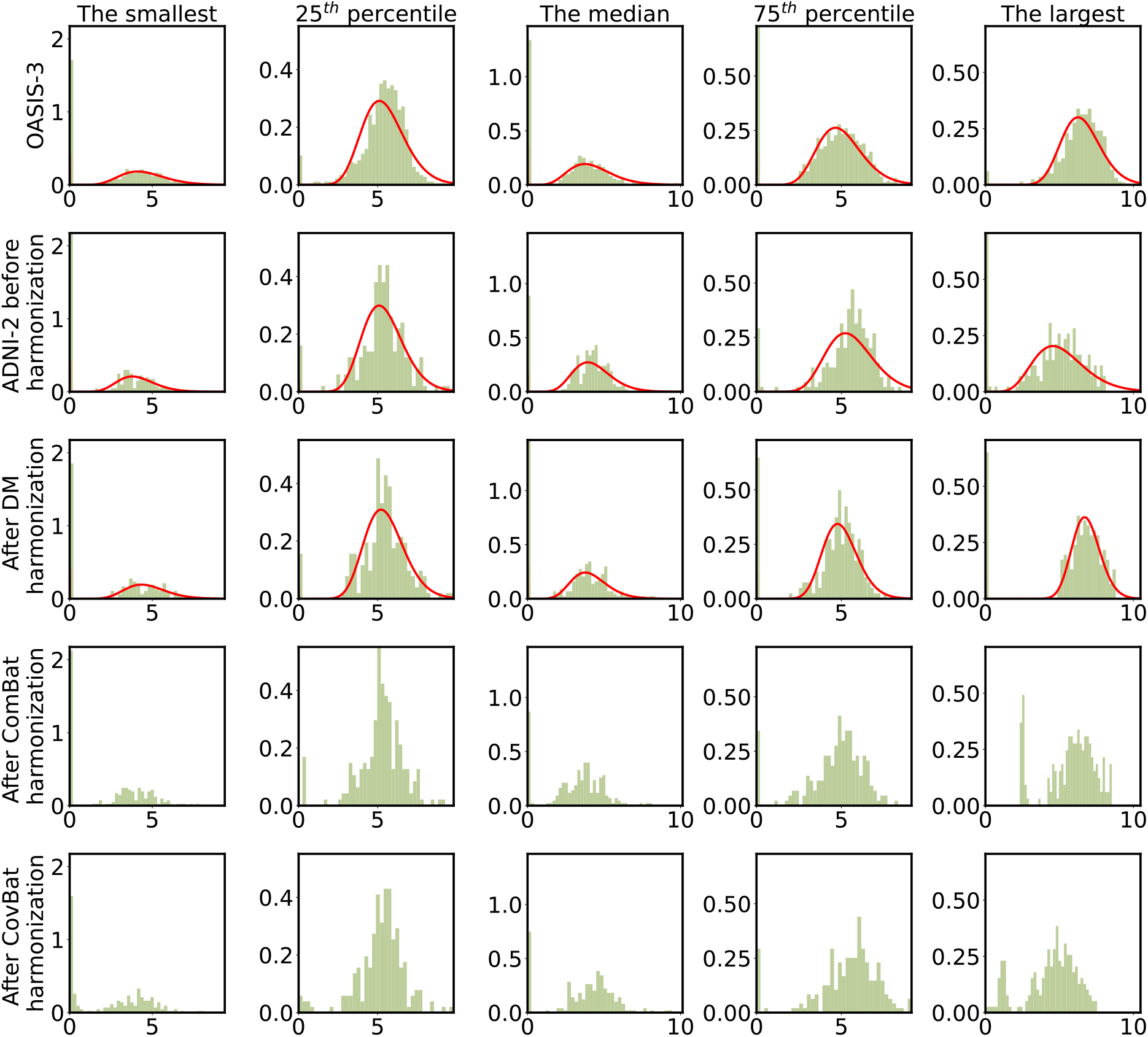
Normalized histograms of five connectivity values selected by choosing the smallest, 25^th^ percentile, median, 75^th^ percentile, and largest improvement in significance values (Δ*s*) from the correlation between MMSE and brain connectivity. Each column represents a structural brain connection, with the first row showing the reference OASIS-3 data and the second row representing the ADNI-2 data before harmonization. The third, fourth, and fifth rows show ADNI-2 data after harmonization using the DM, ComBat, and CovBat methods (all without including covariates), respectively. The red curves show the fitted gamma distribution for OASIS-3, as well as for ADNI-2 before and after DM harmonization. ComBat and CovBat harmonized data display more irregular, multimodal distributions, with some zero values transformed to non-zero values. We projected the negative values produced by ComBat and CovBat to zero for visualization consistency.

As illustrated in Fig. 1, our DM approach achieves effective alignment of connectivity distributions between ADNI-2 and OASIS-3 while preserving the inherent variability across subjects. The method preserves the relative rank ordering of connectivity values within each cohort while adjusting the overall distribution parameters to match between datasets. This preservation of inter-subject variability is essential for detecting true biological relationships. The strengthened correlations with MMSE and age after harmonization (as shown in Section 3.2) further support the conclusion that our method preserves meaningful variability across subjects. Figure 1 also demonstrates the appropriateness of the gamma distribution for modeling positive structural brain connectivity values. The fitted gamma distributions (red curves) closely follow the empirical histograms of positive connectivity values (green bars) across both datasets, capturing both the peak and tail characteristics of the empirical distributions. This visual evidence confirms that our choice of a gamma+delta distribution model is suitable for representing the statistical properties of structural connectivity data in both datasets, providing a solid foundation for our DM harmonization approach.

### 3.2 Correlation between harmonized structural connectivity and MMSE/age

We present the results of the quantitative evaluation of the proposed method in this section and in Tables 2, 3, 4 and Fig. 2. Table 2 presents the correlations between MMSE and connectivity values within each dataset separately, as well as in the combined dataset before and after harmonization. The comparison involves multiple harmonization approaches: the proposed DM (both with and without separating sex groups) and ComBat/CovBat (with and without sex as a covariate).

**Table 2:**
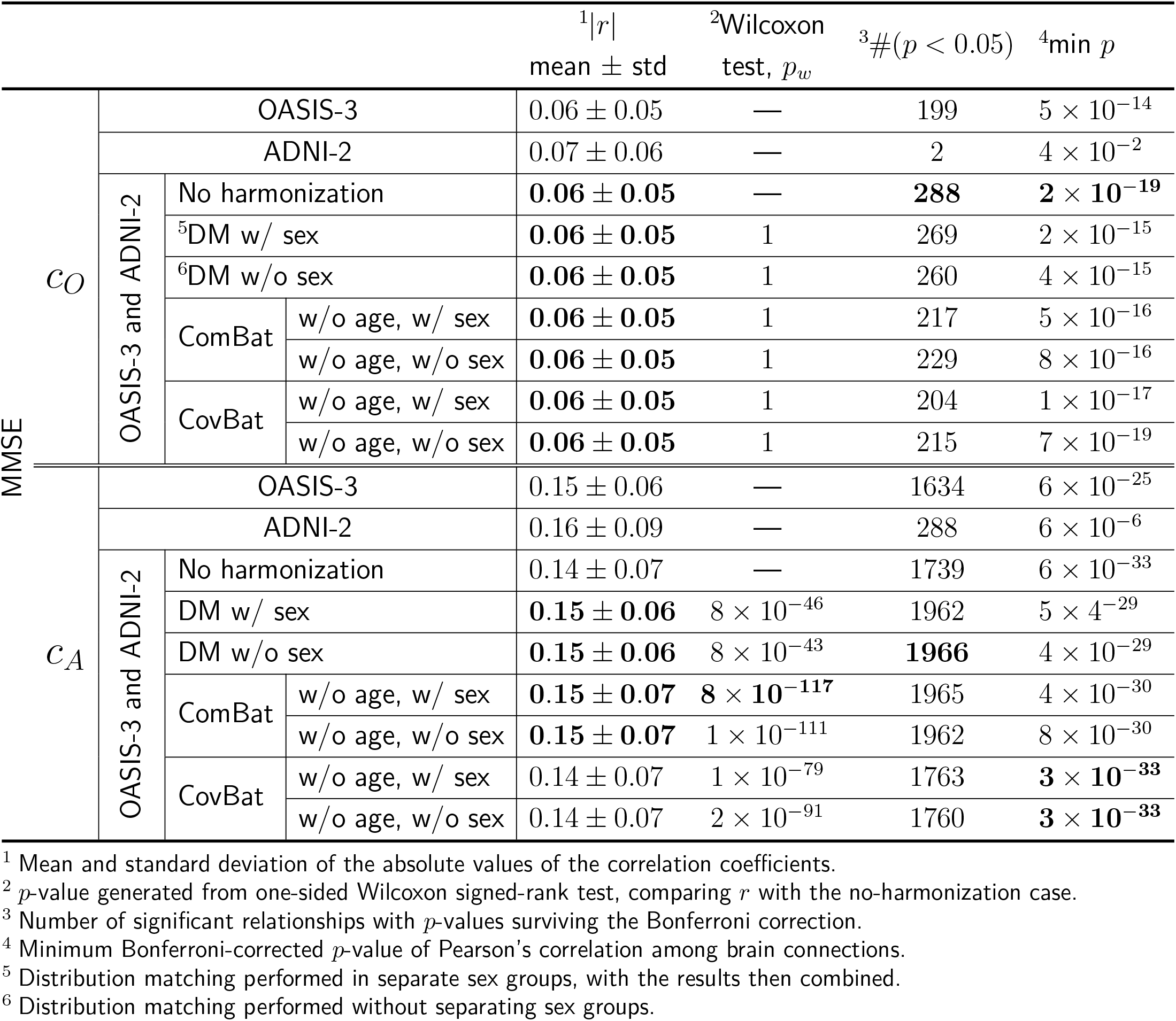
The comparison of the correlations between MMSE and connectivity values before/after harmonization for the ADNI-2 and OASIS-3 datasets.

**Table 3:**
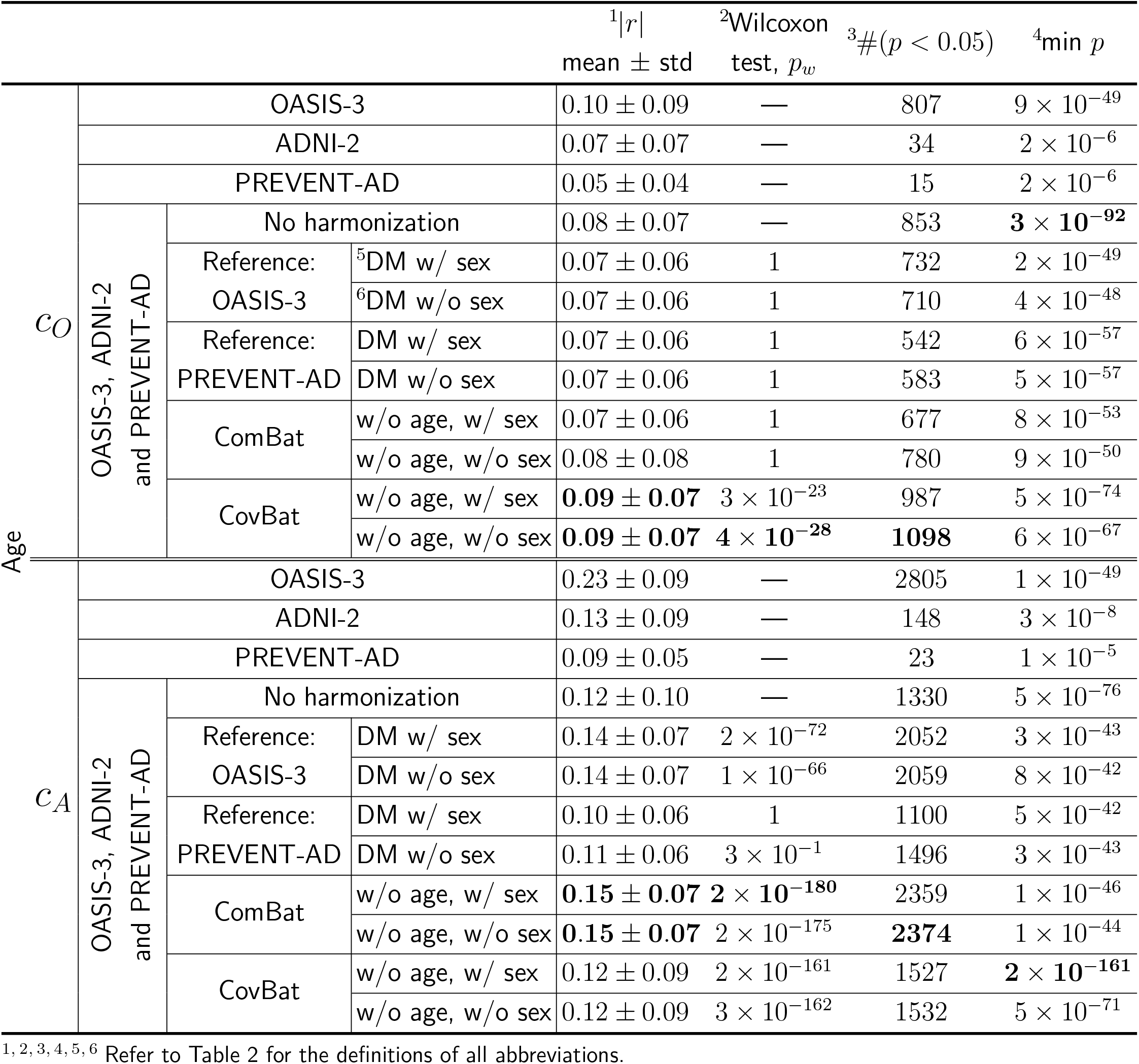
The comparison of the correlations between age and connectivity values before/after harmonization for OASIS-3, ADNI-2, and PREVENT-AD datasets.

**Table 4:**
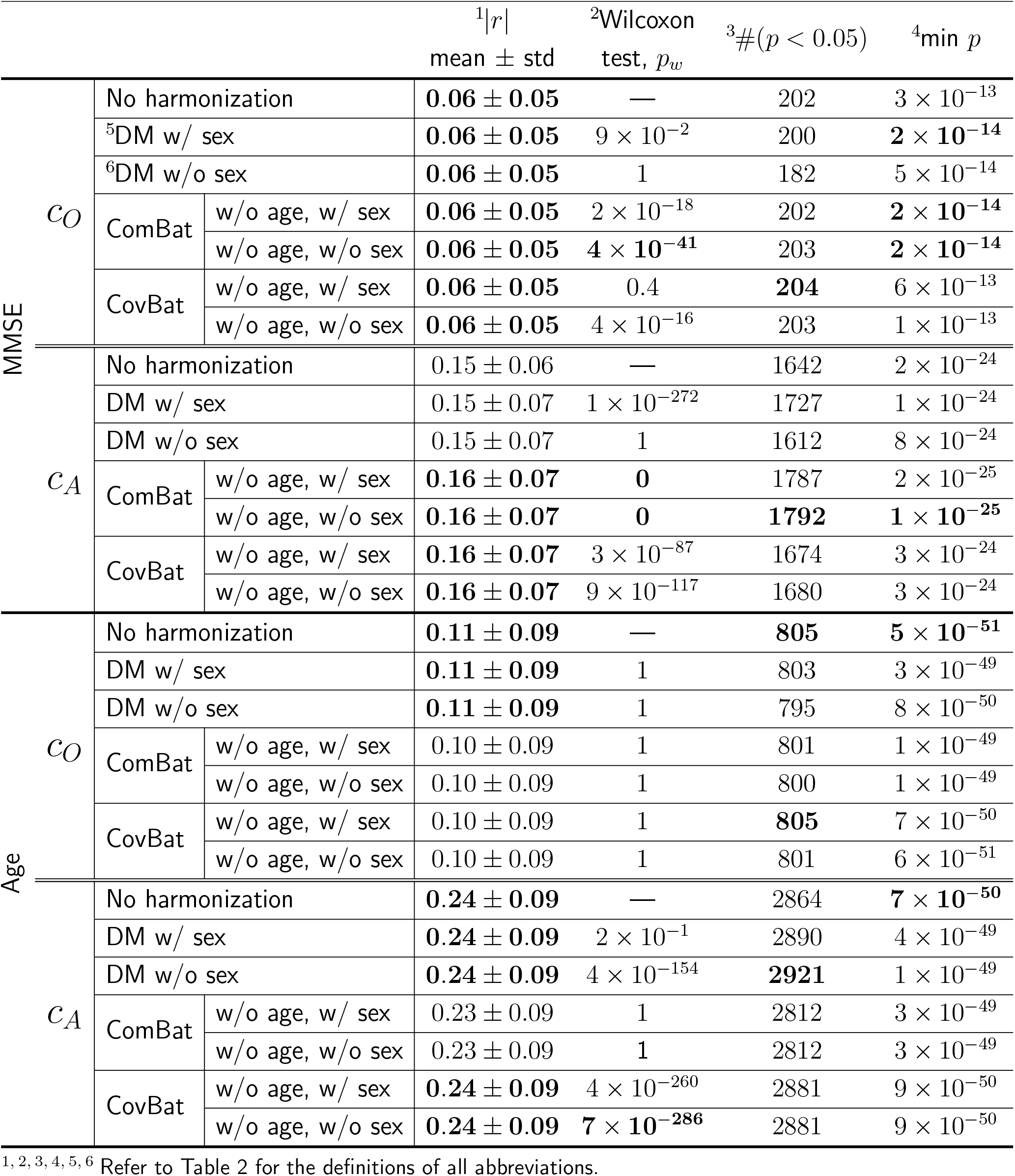
The comparison of the correlations between age/MMSE and connectivity values before/after harmonization for internal harmonization within OASIS-3 dataset.

**Figure 2:**
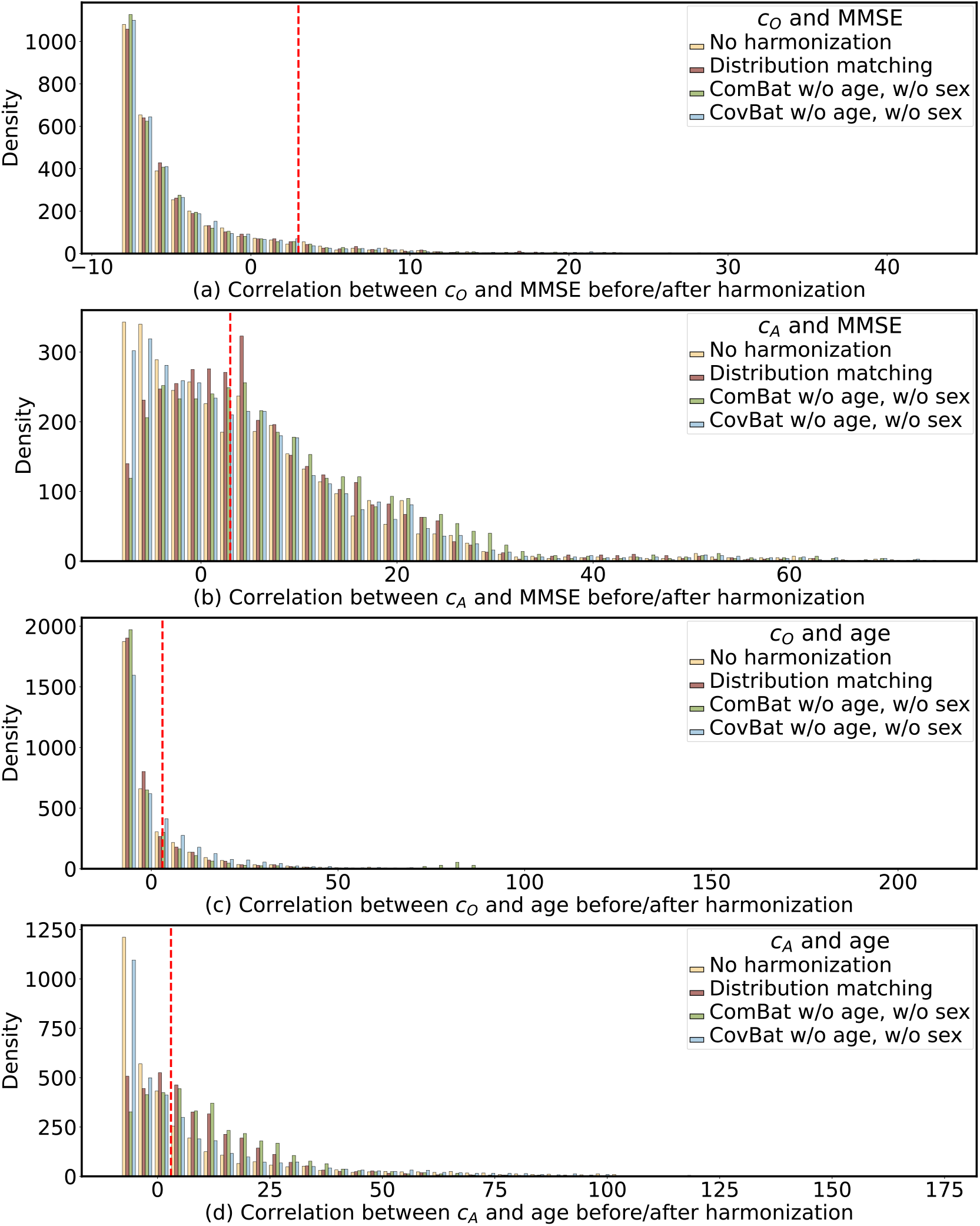
The histogram of *s* := − log *p* from Pearson’s correlation between original (a,c) or augmented (b,d) structural brain connectivity and MMSE (a,b) or age (c,d), before and after harmonization. The red dashed vertical line represents the significance cutoff threshold of − log 0.05.

For the correlation between MMSE and original structural connectivity, DM (both variants) maintained the baseline mean absolute correlation (0.06 *±* 0.05) while identifying a high number of significant relationships (260-269, Table 2, Fig. 2(a)). All variations of ComBat and CovBat showed comparable performance, with similar correlations but fewer significant relationships. In correlations with augmented structural connectivity, DM methods performed well with a correlation strength of (0.15*±*0.06) and 19621966 significant relationships (also shown in Fig. 2(b)), comparable to ComBat’s performance. ComBat variants showed similar results, while CovBat configurations demonstrated slightly lower correlation values and fewer significant relationships but maintained strong statistical significance with smallest minimum *p*-values.

For MMSE correlations with augmented structural connectivity, the DM method showed significant improvement based on Wilcoxon test *p*-values. We expected a positive correlation between MMSE and brain connectivity because lower MMSE indicates more advanced cognitive decline. Figure 3 (top row) demonstrates that DM enhanced the correlation between MMSE and (both original and augmented) structural connectivity, as visible in most dots being above the identity line.

**Figure 3:**
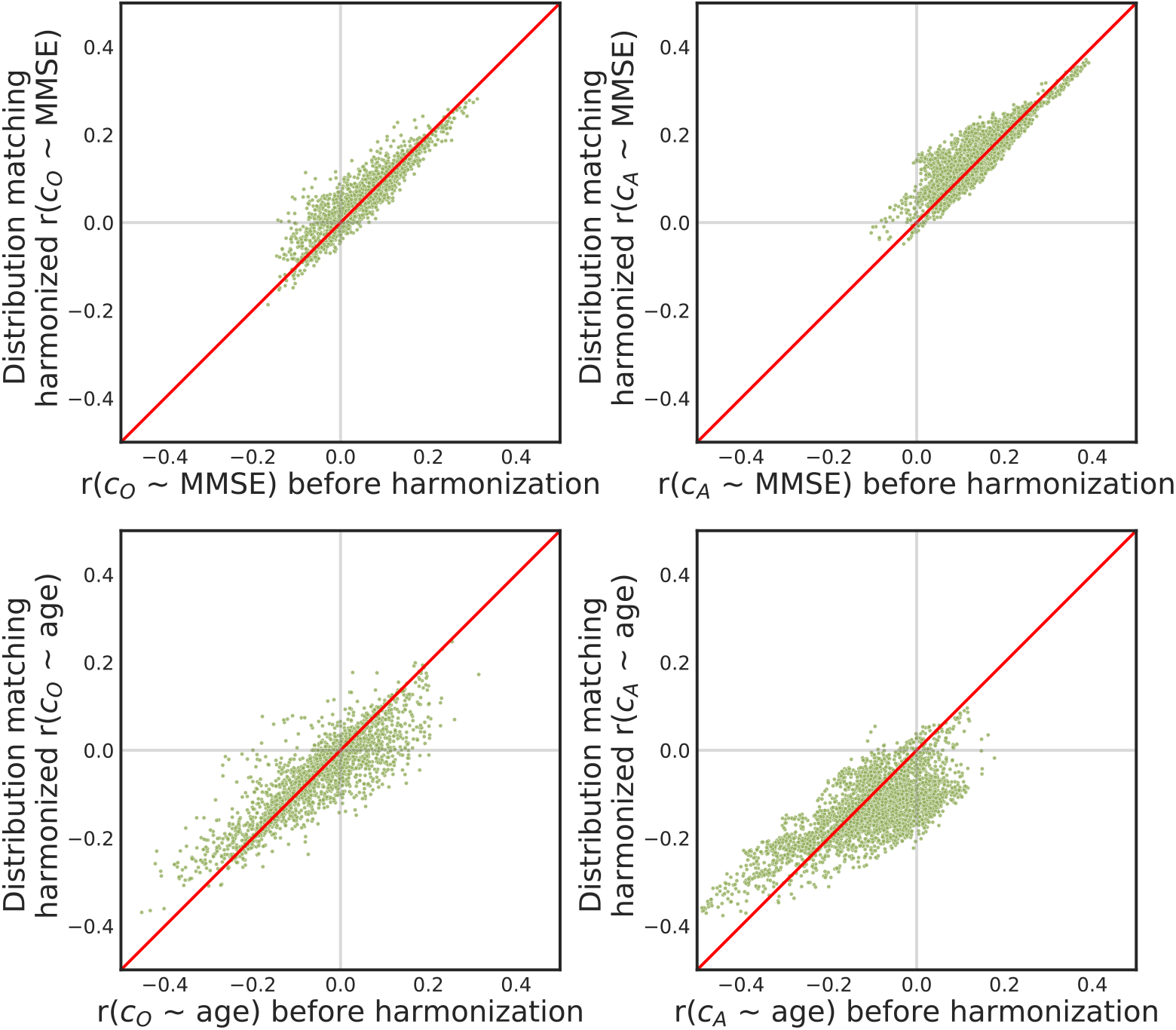
Scatter plots comparing post- vs. pre-harmonization Pearson’s correlation (*r*) between MMSE (top) or age (bottom) and original (left) or augmented (right) structural brain connectivity. The identity line is plotted in red. In all cases, the proposed DM harmonization increased the overall |*r*|.

Table 3 presents the correlations between age and connectivity values across all three datasets (OASIS-3, ADNI-2, and PREVENT-AD) before and after harmonization. The performance of DM varied depending on the chosen reference site, with OASIS-3 as reference generally yielding better results than PREVENT-AD. For original structural connectivity, CovBat demonstrated the strongest age-connectivity correlations (0.09 *±* 0.07) with the highest number of significant correlations (also in Fig. 2(c)). For augmented structural connectivity, our DM approach yielded strong results (0.14 *±* 0.07) with over 2000 significant correlations (also in Fig. 2(d)), although ComBat showed slightly higher correlation values with more significant relationships.

In the case of age, we expected a negative correlation with structural brain connectivity. Figure 3 (bottom row) illustrates a post-DM improvement in the correlation between age and (both original and augmented) structural connectivity, as evidenced by most of the dots being below the identity line.

Next, since the OASIS-3 dataset includes data from two types of Siemens MRI scanners (as shown in Table 1), we performed internal harmonization within that dataset, the results of which are shown in Table 4. Different harmonization methods resulted in only small differences in the mean absolute correlations and the number of significant relationships. In particular, for correlations with age, no significant improvements were observed in the mean absolute correlations except for the correlation with augmented connectivity after DM w/o sex and CovBat methods, all showing similar mean absolute correlations. For MMSE correlations with augmented structural connectivity, ComBat and CovBat methods slightly improved the mean absolute correlations, more than DM did. However, with original structural connectivity, no sizable improvements were observed in the mean absolute correlations of any of the methods.

These comprehensive analyses across different datasets and metrics demonstrate that our DM approach performs well for harmonizing structural brain connectivity, particularly when using appropriate reference sites. As evident in the tables, the correlations were consistently stronger with the augmented than original connectivity. The choice of reference site (OASIS-3 vs. PREVENT-AD) showed notable differences in performance, especially for age correlations with augmented connectivity, suggesting that reference site selection is an important consideration in harmonization studies. Interestingly, performing DM with or without separating sex groups did not lead to substantial differences in results in our experiments, suggesting that our approach demonstrates robustness to this biological variable within the specific datasets examined in this study.

Lastly, regarding the permutation experiments within ADNI-2 (Section 2.5), the PDFs of the 1000 permutations of the subsampled datasets and the full dataset are visualized in Fig. 4, illustrating the extent of variability introduced by cohort size reduction.

**Figure 4:**
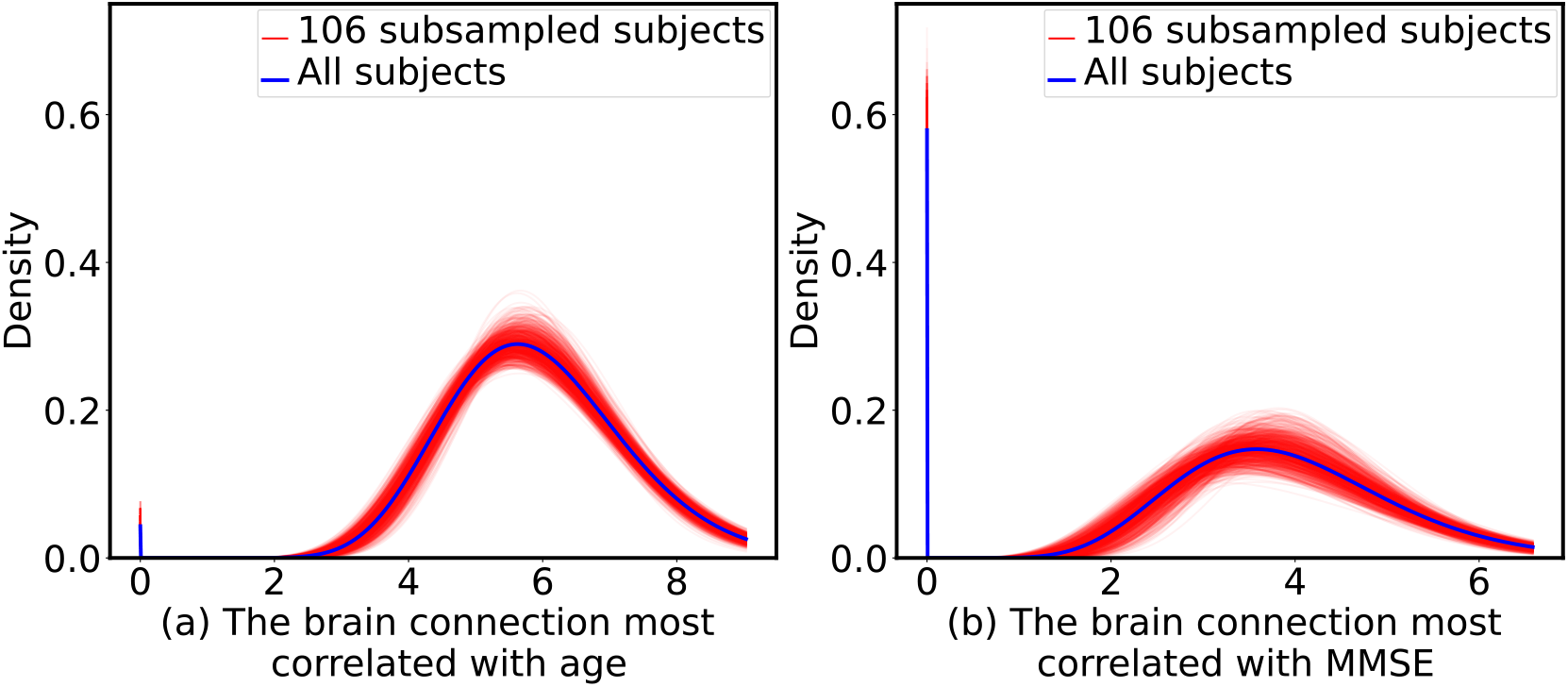
Distribution of the PDFs of the brain connection most correlated with (a) age and (b) MMSE. Each plot compares 1000 random permutations to select a subset of 106 subjects (red curves), to the full dataset with 209 subjects (blue curve), in ADNI-2. The brain connection most correlated with age (a) is between the left thalamus and left hippocampus. The connection most correlated with MMSE (b) is between the left lingual cortex and left middle temporal cortex.

## 4 Discussion

In this study, we proposed a DM technique for harmonizing multi-site structural brain connectivity data, and evaluated it while comparing it with the ComBat and CovBat harmonization approaches. We analyzed structural brain connectivity data from three datasets (OASIS-3, ADNI-2, and PREVENT-AD), and examined the impact of harmonization on how brain connectivity correlates with MMSE and age. We evaluated the efficacy of our DM method both qualitatively and quantitatively, with assessments showing competitive performance across different metrics. Qualitatively, our DM harmonization achieved the expected distributional alignment across datasets, as demonstrated in Fig. 1. The effectiveness of harmonization was assessed quantitatively by comparing correlation strengths and statistical significance after applying each harmonization method to those before harmonization. We also examined the impact of reference-site selection by using both OASIS-3 and PREVENT-AD as reference sites in separate experiments, and found that reference-site choice significantly affected harmonization outcomes. The option to process sex groups separately or together provides additional flexibility to account for biological differences while removing site-specific effects. Through our cross-dataset analyses, we found DM produced comparable or sometimes better results than alternative methods did in many scenarios (Tables 2 and 3). Our findings provide insight into the complexities of harmonizing multi-site structural brain connectivity data and lay the groundwork for further refinements in harmonization techniques.

Our analysis intriguingly revealed that using different reference sites for harmonization could substantially impact the enhancement of biological correlations. We found that choosing OASIS-3 (761 subjects) as the reference site resulted in stronger connectivity-age correlation than choosing PREVENT-AD (340 subjects) as the reference site did. This is consistent with the intuitive preference towards choosing the larger dataset as reference, as it would minimize the amount of data transferred and maximize the amount of data remaining intact.

While minimum *p*-values can provide insight into the strongest individual correlations, relying solely on this measure has some limitations in the context of large-scale neuroimaging studies. In datasets involving thousands of brain connections, extremely low *p*-values can occur by chance, potentially leading to misleading conclusions about harmonization effectiveness. Moreover, minimum *p*-values are not robust to outliers or extreme values, which could disproportionately affect the final conclusions. To address these limitations and provide a more comprehensive assessment of harmonization performance, we incorporated the following additional measures. Absolute correlation coefficients offer a direct indication of the strength of association between brain connectivity and MMSE/age, providing insight into effect sizes that are theoretically independent of sample size and reflect practical significance. Wilcoxon signed-rank tests, comparing results before and after applying different harmonization methods, help us understand whether harmonization significantly improves the overall strength of correlations across all connections, providing a whole-brain measure of effectiveness. (This test requires the conditional distribution of each observation given the others to be symmetric about a common point, which may, however, not always be satisfied in our case, hence reduced accuracy of the Wilcoxon *p*-values.) The number of connections surviving Bonferroni correction reflects the overall sensitivity for detecting meaningful correlations while controlling for multiple comparisons, providing insight into the breadth of harmonization effects throughout all brain connections. By utilizing these multiple measures, we aimed to capture different aspects of harmonization performance. This could offer a more comprehensive and nuanced evaluation that enables the assessment of both the strength and prevalence of brain-behavior relationships across the entire brain, providing a more reliable basis for comparing different harmonization techniques.

Our evaluation consisted of DM harmonization on three datasets combined, as well as internal harmonization within a single (OASIS-3) dataset. Interestingly, we observed that internal harmonization within OASIS-3 rarely improved the correlation results, regardless of whether our DM, ComBat, or CovBat was used (see Table 3). This outcome could be attributed to the following two factors. The OASIS-3 dataset has a high degree of consistency and standardization, e.g. in terms of data collection protocols, acquisition parameters, and possibly some preprocessing. If there is minimal variability between different scanner types or parameters within OASIS-3, the impact of harmonization will be negligible, thereby making internal harmonization less beneficial. Furthermore, in the harmonization of the three datasets combined, all datasets had at least 209 subjects, whereas in the internal harmonization within OASIS-3, we divided OASIS-3 into two sites: one with 655 subjects (used as the reference site) and the other with 106 subjects (used as the new site). The reduced number of subjects in the new site (from 209 to 106) could be a contributing factor. We performed permutation tests by randomly selecting 106 subjects from the total 209 subjects in the ADNI-2 dataset to simulate the reduction in the sample size in our internal harmonization experiment, and computed the distribution of the selected feature. As shown in Figure 4, we found a non-negligible variation among the permuted PDFs (red curves), which often deviated from the full sample distribution (blue curve) to some extent. This suggests that a smaller new site might represent the variability and distribution less precisely, leading to less robust harmonization. The lack of improvement in correlation results within the OASIS-3 dataset after internal harmonization could therefore be attributed to the possible homogeneity of the dataset and the smaller new site in internal harmonization. Accounting for these factors can help us refine harmonization techniques and better tailor them to specific datasets, ultimately improving the reliability and consistency of large-scale multi-site neuroimaging analyses.

We compared the DM method with two variants of ComBat and CovBat harmonization methods on structural connectivity data, one using only sex as a covariate and the other without covariates. The inclusion of demographic covariates in harmonization aims to remove site-specific variability attributable to these factors, potentially harmonizing data more effectively across different sites, age ranges, and sex groups (Pomponio et al., 2020; Zhou et al., 2023). Yet this approach inherently creates connectivity measurements that are partly dependent on these demographic variables. The fundamental challenge in harmonization lies in distinguishing between site-specific technical artifacts and genuine biological variations. While incorporating demographic covariates, like age, seems to protect biological signals, it also risks creating analysis bias when these variables become outcome measures (An et al., 2022). To address this methodological issue, our comparative analyses excluded ComBat/CovBat configurations containing age covariates when evaluating connectivity relationships. Our DM approach circumvents this pitfall by operating independently of explicit covariate modeling, yet achieves effective harmonization while preserving biological associations with outcome variables. Nonetheless, we have demonstrated the flexibility of our approach through the results by a variation of our DM method that incorporated sex as a covariate. If there is a mismatch between the distributions of a *continuous* biological variable, *x* (e.g. age), among sites, and provided that sample sizes are sufficient, the effect can be removed by subsampling the data to achieve similar distributions of *x* between sites prior to harmonization. Otherwise, our DM approach can nevertheless be extended to control for such continuous biological variables. For instance, the strength of a brain connection for subject *i* of a certain site can be modeled as 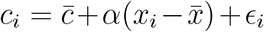, where 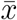 is the within-site average of *x, ϵ*_*i*_ is the redisual error, and the constants 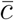and *α* are fitted through linear regression. We can subsequently apply our DM method to 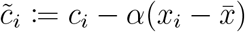, which should no longer carry the (linear) effect of *x*. We do not, however, suggest taking this regression approach if *x* (or a variable correlated with *x*) is used in the downstream analysis, given the abovementioned information leakage. That is why we did not control for age in our age or MMSE correlation analysis (note the strong correlation between age and MMSE in Section 2.1).

Another important advantage of our method, as opposed to ComBat/CovBat, is its preservation of the structural integrity of the brain connectivity matrices, specifically the nonnegativity of connectivity and the maintenance of zero values in connectivity matrices. The sparsity of (our original) structural connectivity reflects the absence of direct anatomical pathways between many regions; transforming these zeros into non-zero values, as can occur with ComBat/CovBat (Fig. 1), complicates biological interpretation. Our method is specifically designed to maintain zeros as zeros throughout the harmonization process, ensuring that the physical interpretation of connectivity remains consistent. This preservation of sparsity patterns is crucial for accurate representation of brain network topology. While a non-zero-preserving variant (with *u*(0) = 1 in Eq. (5)) might yield higher correlation scores (as briefly tested), we prioritize the interpretational consistency afforded by the primary DM design. Nonetheless, we have demonstrated the flexibility of our approach through the results of a variation of our DM method that incorporated sex as a covariate, while still maintaining the zero-preservation property.

As shown in Fig. 1, we selected five brain connections by ranking the difference of negative logarithm of *p*-values (Δ*s*) generated from the correlation between MMSE and structural brain connectivity before vs. after harmonization, and choosing the smallest, 25^*th*^ percentile, median, 75^*th*^ percentile, and largest ones. We chose these five examples in this way so they were selected in an unbiased manner. This, however, might create the expectation that the harmonization-caused shift in the peak of the histogram would increase with respect to Δ*s*. However, as shown in the figure, the amount of change after harmonization does not necessarily correlate with the ranking of Δ*s*. It is worth noting that Δ*s* is related to not only the harmonization effect size, but likely also to the significance (*s*) of the correlation with MMSE itself, with the latter varying largely across different structural brain connections and not expected to be necessarily related to the effectiveness of harmonization. In other words, brain connections that benefit the most from harmonization and possibly go through a sizable distribution shift may not necessarily coincide with those that are highly correlated with MMSE and have a high Δ*s*.

Our current study has several limitations that present opportunities for future research. First, our analysis was confined to three imaging datasets and one tractography method, which limits the generalizability of our findings. Future studies should extend this work to include additional datasets from diverse sites and data processed with various tractography methods, allowing for a more comprehensive evaluation of the DM method. This could include experiments with datasets from different age ranges, such as children or young adults, to test the method’s effectiveness across the lifespan. Second, while we have used age and MMSE as correlation variables in our current analysis, a limitation is that our experimental results do not precisely quantify how much specific effects (e.g., scanner effects) were reduced by harmonization. While the main motivation of harmonization is to reduce site-dependent hardware and acquisition effects, further work is needed to disentangle and quantify the specific contributions of harmonization to reducing various non-biological sources of variation. Additional demographic and clinical variables could be included in future analyses to address this limitation. Third, to further validate the effectiveness of our harmonization approach, downstream analyses, such as predictive modeling, classification, and more complex network analyses could be employed. Such analyses would provide concrete evidence of the impact of harmonization on other applications in neuroimaging research, potentially revealing subtle effects that might be obscured by harmonization methods less tailored to structural brain connectivity. Specifically, prediction tasks (e.g., age prediction and diagnosis classification) could provide another validation dimension beyond correlation analysis, though such tasks would introduce additional variables related to machine learning algorithms, model architectures, and training protocols that could confound the assessment of the harmonization method itself. Finally, future work could investigate the impact of harmonization on longitudinal data, explore its effectiveness in harmonizing different imaging modalities, and examine its performance in clinical populations with specific neurological or psychiatric conditions. Addressing these limitations and exploring these new directions, which would enhance the robustness, applicability, and sensitivity of our DM harmonization in multi-site neuroimaging studies, is part of our ongoing research.

## Data and Code Availability

Data used in this study were obtained from three public databases: 1) the third phase of the Open Access Series of Imaging Studies (OASIS-3) (LaMontagne et al., 2019), which is freely available (www.oasis-brains.org); 2) the second phase of the Alzheimer’s Disease Neuroimaging Initiative (ADNI-2), which is available for download (http://adni.loni.usc.edu) for researchers who meet the criteria for access to these data; 3) the Pre-symptomatic Evaluation of Experimental or Novel Treatments for Alzheimer’s Disease (PREVENT-AD), which can also be freely downloaded (https://openpreventad.loris.ca/).

The toolbox for reconstructing the diffusion orientation distribution function in constant solid angle, performing Hough-transform global probabilistic tractography, computing the connectivity matrix, and augmenting it with indirect connections is available at www.nitrc.org/projects/csaodf-hough. ComBat data harmonization was performed using a package available at https://github.com/rpomponio/neuroHarmonize. FreeSurfer (Fischl, 2012) and FSL (Jenkinson et al., 2012) are available for download at https://freesurfer.net and https://fsl.fmrib.ox.ac.uk, respectively. All other code implementing our distribution-matching approach is available at https://github.com/zzstefan/Distribution_matching.

## Author Contributions

Zhen Zhou: Conceptualization, Methodology, Formal analysis, Validation, Writing - original draft, and Writing - review & editing. Bruce Fischl: Conceptualization, Methodology, and Writing - review & editing. Iman Aganj: Data curation, Conceptualization, Methodology, Formal analysis, Writing - review & editing, and Funding acquisition.

## Funding

Support for this research was provided by the Michael J. Fox Foundation for Parkinson’s Research (MRI Biomarkers Program award MJFF-021226), as well as in part by the National Institutes of Health (NIH), specifically the National Institute on Aging (NIA; RF1AG068261, R01AG068261).

Additional support was provided by the NIH, specifically in part by the NIA (R21AG082082, R01AG064027, R01AG008122, R01AG016495, R01AG070988), the BRAIN Initiative Cell Census Network grants U01MH117023 and UM1MH130981, the Brain Initiative Brain Connects consortium (U01NS132181, UM1NS132358), the National Institute for Biomedical Imaging and Bioengineering (R01EB023281, R01EB006758, R21EB018907, R01EB019956, P41EB030006), the National Institute of Mental Health (UM1MH130981, R01MH123195, R01MH121885, RF1MH123195), the National Institute for Neurological Disorders and Stroke (R01NS0525851, R21NS072652, R01NS070963, R01NS083534, U01NS086625, U24NS10059103, R01NS105820), and was made possible by the resources provided by Shared Instrumentation Grants S10RR023401, S10RR019307, and S10RR023043. Additional support was provided by the NIH Blueprint for Neuroscience Research (U01MH093765), part of the multi-institutional Human Connectome Project.

Data collection and sharing for this project was funded in part by OASIS-3: Longitudinal Multimodal Neuroimaging: Principal Investigators: T. Benzinger, D. Marcus, J. Morris; NIH P30 AG066444, P50 AG00561, P30 NS09857781, P01 AG026276, P01 AG003991, R01 AG043434, UL1 TR000448, R01 EB009352.

Data were also provided in part by the Alzheimer’s Disease Neuroimaging Initiative (ADNI) (NIH Grant U01 AG024904) and DOD ADNI (Department of Defense award number W81XWH-12-2-0012). ADNI is funded by the NIA, the NIBIB, and through generous contributions from the following: AbbVie, Alzheimer’s Association; Alzheimer’s Drug Discovery Foundation; Araclon Biotech; BioClinica, Inc.; Biogen; Bristol-Myers Squibb Company; CereSpir, Inc.; Cogstate; Eisai Inc.; Elan Pharmaceuticals, Inc.; Eli Lilly and Company; EuroImmun; F. Hoffmann-La Roche Ltd and its affiliated company Genentech, Inc.; Fujirebio; GE Healthcare; IXICO Ltd.; Janssen Alzheimer Immunotherapy Research & Development, LLC.; Johnson & Johnson Pharmaceutical Research & Development LLC.; Lumosity; Lundbeck; Merck & Co., Inc.; Meso Scale Diagnostics, LLC.; NeuroRx Research; Neurotrack Technologies; Novartis Pharmaceuticals Corporation; Pfizer Inc.; Piramal Imaging; Servier; Takeda Pharmaceutical Company; and Transition Therapeutics. The Canadian Institutes of Health Research is providing funds to support ADNI clinical sites in Canada. Private sector contributions are facilitated by the Foundation for the NIH (www.fnih.org). The grantee organization is the Northern California Institute for Research and Education, and the study is coordinated by the Alzheimer’s Therapeutic Research Institute at the University of Southern California. ADNI data are disseminated by the Laboratory for Neuro Imaging at the University of Southern California.

Data were also obtained in part from the Pre-symptomatic Evaluation of Novel or Experimental Treatments for Alzheimer’s Disease (PREVENT-AD) program.

## Acknowledgments

We thank Dr. Aina Frau-Pascual for previously helping us in data curation.

